# Parallel networks to predict TIMP and protease cell activity of Nucleus Pulposus cells exposed and not exposed to pro-inflammatory cytokines

**DOI:** 10.1101/2024.08.28.609099

**Authors:** L. Baumgartner, S. Witta, J. Noailly

## Abstract

**Background:** Intervertebral disc (IVD) degeneration is characterized by a disruption of the balance between anabolic and catabolic cellular processes. Within the Nucleus Pulposus (NP), this involves increased levels of the pro-inflammatory cytokines Interleukin 1beta (IL1B) and Tumor Necrosis Factor (TNF) and an upregulation of the protease families MMP and ADAMTS. Primary inhibitors of those proteases are the tissue inhibitors of matrix metalloproteinases (TIMP). This work aims at contributing to a better understanding of the dynamics among proteases, TIMP and proinflammatory cytokines within the complex, multifactorial environment of the NP.

**Methods:** The Parallel Network (PN)-Methodology was used to estimate relative mRNA expressions of TIMP1-3, MMP3 and ADAMTS4 for five simulated human activities; walking, sitting, jogging, hiking with 20 kg extra weight, and exposure to high vibration. Simulations were executed for nutrient conditions in non- and early-degenerated IVD approximations. To estimate the impact of cytokines, the PN-Methodology inferred relative protein levels for IL1B and TNF, re-integrated as secondary stimuli into the network.

**Results:** TIMP1 and TIMP2 expression were found to be overall lower than TIMP3 exp. In absence of pro-inflammatory cytokines, MMP3 and/or ADAMTS4 expression were strongly downregulated in all conditions but vibration and hiking with extra weight. Pro-inflammatory cytokine exposure resulted in an impaired inhibition of MMP3, rather than of ADAMTS4, progressively rising with increasing nutrient deprivation. TNF mRNA was less expressed than IL1B. However, at the protein level, TNF was mainly responsible for the catabolic shift in the simulated pro-inflammatory environment. Overall, results agreed with previous experimental findings.

**Conclusions:** The PN-Methodology successfully allowed the exploration of the relative dynamics of TIMP and protease regulations in different mechanical, nutritional, and inflammatory environments, in the NP. It shall stand for a comprehensive tool to integrate in vitro model results in IVD research and approximate NP cell activities in complex multifactorial environments.

## 1 Introduction

Low back pain is the leading cause of disability worldwide, presenting both an economic and epidemiological burden [1]. It affects 60-80% of people globally, predominantly individuals in middle to old age imposing severe restrictions in social and physical capacity and importantly affecting mental health [2,3].

The onset of low back pain is strongly correlated with intervertebral disc (IVD) degeneration (IDD) [4]. IDD is characterized by a disruption of the balance between catabolic and anabolic protein secretion, which increases the presence of proteases that degrade the IVD extracellular matrix [5–10]. The main inhibitors of proteases are the tissue inhibitor of matrix metalloproteinases (TIMP). Although the inhibitory activity of TIMP is recognized, its effective control and role to maintain the integrity of the IVD in a multifactorial environment remains poorly apprehended.

In addition to an increase in protease activity, pro-inflammatory cytokine expression increases during IDD. Especially interleukin 1 beta (IL1B) and tumor necrosis factor (TNF) have been shown to be elevated in degenerated discs [11,12]. IL1B treatment in non-degenerated Nucleus Pulposus (NP) cells led to upregulated gene expression of key catabolic proteins including MMP3 and ADAMTS4 [5]. Elevated TNF levels were associated with non-recoverable catabolic shifts in the IVD [13]. However, the role of proinflammatory cytokines in the dual regulation of proteases and TIMP has not yet been elucidated.

According to experimental models, nutritional and mechanical extrinsic stimulations are believed to largely influence IVD cells [10]. As the IVD is the largest avascular structure within the human body [14]cell nutrition depends on the diffusion of nutrients from the capillaries located at the Cartilage Endplates [15]. This creates steep gradients of nutrients (oxygen, glucose, pH), resulting in low glucose and oxygen concentrations and low pH around the mid transverse-plane of the NP [15]. Additionally, human movement activities cause the IVD to be dynamically exposed to a variety of mechanical loads, translated into varying intradiscal pressures (magnitude) and frequencies into the NP [16]. Despite extensive studies evaluating the effect of key nutritional (pH, glucose) and mechanical (magnitude, frequency) stimuli on gene expression of TIMP and proteases [9,17–20], insight on the combined effect of regulatory stimuli on NP cell activity, in absence or presence of pro-inflammatory cytokines is still limited. Arguably, experimental research remains time and cost intensive, and straightforward in silico methods to approximate dynamics at a (multi-)cellular level remain scarce.

To cope with this, a methodology was recently proposed to simulate cell activity (CA) regulated by multifactorial stimulus environments, which was mathematically represented as simultaneous ongoing actions in parallel networks (PN) [21]. This method combines experimental data about cell responses to varying stimulus doses, therefore, allowing to estimate effective CA directly at the (multi-)cellular level, while considering intracellular regulations as a “black box”. The PN-Methodology also allows to automatically switch the nature of a stimulus (i.e. activating or inhibiting), depending on the dose of this stimulus, as informed by experimental models.

Hence, the objective of this work is to investigate the dynamics of TIMP subgroups 1-3, MMP3 and ADAMTS4 in response to complex microenvironmental stimuli, with and without TNF and IL1B proteins. To this end, the PN-Methodology was further leveraged regarding mathematical estimations of mRNA-protein dynamics.

## 2 Materials and Methods

### 2.1 Overview of the simulated network

mRNA expressions of TIMP1, TIMP2, TIMP3, MMP3, ADAMTS4, IL1B and TNF were estimated based on user-defined levels of key nutritional and mechanical stimuli, within ranges of stimulus doses defined in the literature (Figure 1). Stimulus doses were set to simulate five human activities: sitting, walking, hiking with 20kg extra weight and exposure to vibration. Each activity was simulated for three nutritional conditions, covering optimal and early degenerated cell environments.

**Figure 1:**
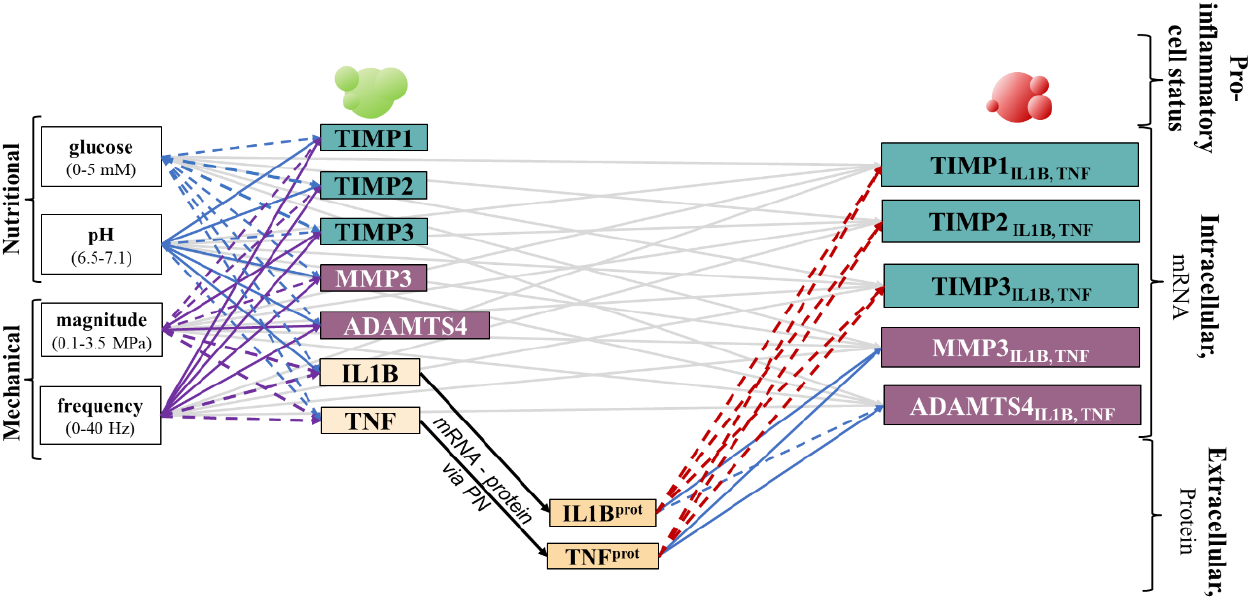
Overview of the simulated environment. The PN-Methodology was used to estimate the relative mRNA expressions of the targeted proteins MMP3, ADAMTS4, TIMP1, TIMP2 and TIMP3 exposed and not exposed to pro-inflammatory cytokines. For cells not exposed to pro-inflammatory cytokines (green colored cells), mRNA expressions were based on glucose concentration, pH, load magnitude, and load frequency, whilst cells exposed to proinflammatory cytokines (red colored cells) additionally obtained input from IL1B and TNF proteins (prot). The nature of the links may be generally activating (blue), generally inhibiting (red), or depending on the stimulus dose (purple). Links reported to be statistically significant in the experimental literature are marked with a continuous line; those reported to be non-significant are reflected by a dashed line. Grey S-CA relationships have the same value than their counterpart in the cytokine-free environment. Extracellular: the PN-methodological concept was used to estimate protein levels of IL1B and TNF, required to estimate a pro-inflammatory environment.

mRNA levels of IL1B and TNF were translated into protein levels, and re-fed into the PN-System as secondary stimuli, to obtain TIMP1, TIMP2, TIMP3, MMP3, ADAMTS4 of cells exposed to a pro-inflammatory environment (Figure 1).

As opposed to nutritional stimuli that define a baseline activation and are, therefore, always activating (see [21] for more information), the nature of mechanical stimuli (magnitude, frequency) depends on the stimulus dose. The effect of pro-inflammatory cytokines were assumed to be either only activating or only inhibiting (Figure 1).

### 2.2. Overview of the Methodology used

The PN-Methodology [21] interprets each CA as an interdependent parallel ongoing action of a cell. Thereby, each CA (here mRNA expression) is determined by combinations of the key nutritional stimuli glucose, pH, magnitude and frequency and reflect one parallel network (PN). Accordingly, many CA are reflected as many relatively small feedforward PN. The relationship between one stimulus with one targeted mRNA expression is called S-CA relationship and is determined by the product of the two variables *θ* and *x*. Both are sensitive to the specific stimulus (subscript S) and cell activity (superscript CA) and therefore specified as 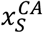 and 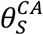, respectively.

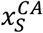 approximates the sensitivity of an mRNA expression to a given stimulus dose through a function. This function is based on discrete values from the literature and assigns a CA to a stimulus dose. If a stimulus is only activating or inhibiting, the CA ranges between 0 (minimal mRNA expression at a given stimulus dose) and 1 (maximal mRNA expression at a given stimulus dose). If the nature of a CA is dose-dependent, it ranges between −1 and 1. Thereby, positive values reflect an activating and negative values an inhibiting effect as previously explained [21] (see Figure 3, results section); the more the CA tends to 1 or to −1, respectively, the higher is the activating or inhibiting effect of the stimulus dose on the CA.

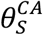 is a (real) number bigger than 0 and maximum 1. It reflects the impact of a stimulus on a given CA (weighting factor). It is derived from the maximal change in mRNA expression, optimally directly calculated from the experimental measurements used to determine 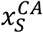. The in-depth pipeline to obtain 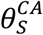 [22] and 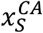 [23] were provided before and visualized in Figure 2.

**Figure 2:**
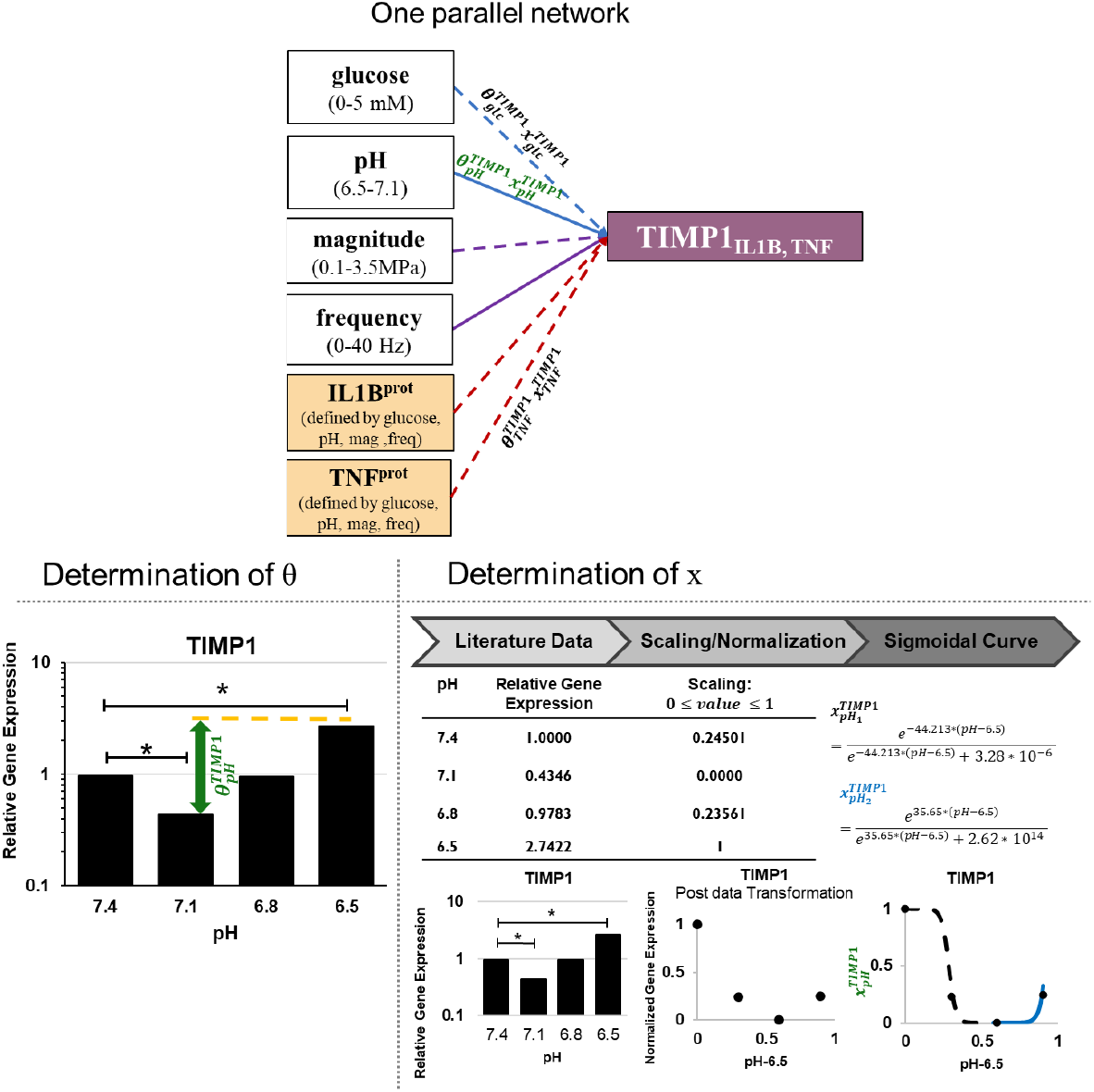
Top: One parallel network of TIMP1 for cells exposed to pro-inflammatory cytokines (see also Figure 1). Stimulus-cell activity (S-CA) relationships are exemplarily mentioned for glucose (glc)-TIMP1, TNF-TIMP1 and pH-TIMP1 (green). Below: Data extraction for pH-TIMP1. Determination of θ: experimental data about TIMP1 mRNA expression at different pH (modified from [17]). θ reflects the highest change in relative gene expression. Determination of x: relative gene expression at pH 7.4, 7.1, 6.8 and 6.5 were normalized to fit continuous sigmoidal functions to approximate a continuous cell activity for TIMP1 mRNA expression throughout a physiological range of pH.

**Figure 3:**
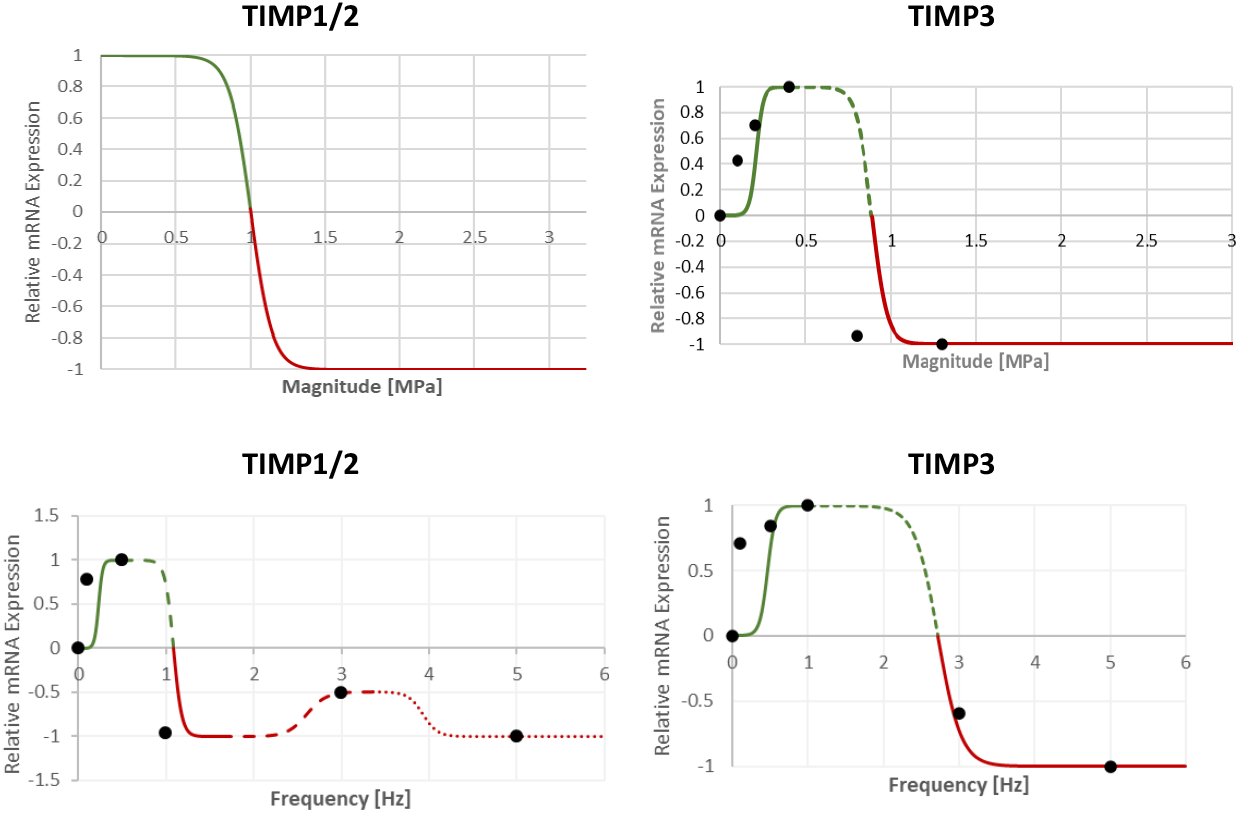
simulated response of TIMP to magnitude and frequency. Due to a lack of data, it was initially assumed that TIMP 2 follows a similar behavior to TIMP1. The relationship of TIMP 1 and TIMP2 to magnitude is based on generic, knowledge-based assumptions [21] Activating effect of loading on TIMP is reflected with values between 0 and 1 (green). Inhibiting effects of loading on TIMP is reflected by values between 0 and −1 (red).

To describe the network given in Figure 1, 38 links (S-CA relationships) need to be specified, each in terms of the sensitivity of a CA to varying stimulus dose 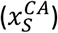 and in terms of the overall impact this stimulus might have on a 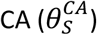.

Whenever possible, CA should be estimated for physiologically relevant stimulus doses. For the NP, glucose concentrations range between 0 mM – 5 mM [24], pH ranges between 6.5 – 7.4 [17] and magnitude varies between 0.1 MPa – 3.5 MPa [16,25]. For frequency, a relevant range for human daily activities was chosen, starting from 0 Hz (static) – 40 Hz to simulate whole body vibrations, such as caused by vehicle engines.

In terms of cell type, human (non-degenerated) cells were prioritized over bovine cells. Otherwise, in vitro work with rat cells was used (Appendix A). 3D cell cultures were preferred over 2D cell cultures. As NP cells change their biological activity as they progress from immature to mature, mature NP cells were preferred.

The activity of each PN was calculated with the PN-Equation (Eq. 1) [21,22] that determines the individual activation and strength of each PN within the PN-System. Briefly: the PN-Equation consists of an activating (α) and an inhibiting (β) portion. The activating portion consists of two terms; in its first term, it considers all the activating weighting factors of the PN-System (*θ*_*α*_). This includes the potentially activating contribution of dose-dependent stimuli. The first activating term is then multiplied with the activating S-CA relationships of the actual 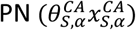. This gives the weight of the tackled PN in the light of the whole PN-System. The inhibiting portion consists of three terms and is calculated independently from the activating term. The first inhibiting term determines the relative strength of the inhibitors for this mRNA. The second term includes all the inhibiting weighting factors for a given mRNA, independently of the CS. The third inhibiting term, eventually, calculates the actual inhibiting S-CA relationship of this PN.

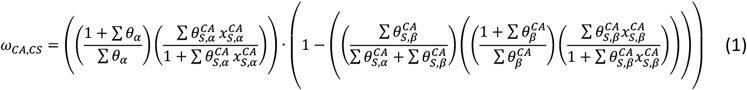

To explicitly assign S-CA relationships to either the activating or inhibiting term of the PN-Equation, and to cope with the dose dependency of magnitude and frequency being either anabolic or catabolic, four sets of PN-Equations were required, where (i) both, magnitude and frequency were anabolic (ii) magnitude was anabolic and frequency catabolic (iii) frequency anabolic and magnitude catabolic and (iv) both, magnitude and frequency were catabolic.

The solution to the PN-Equation provides a dimensionless value called “PN-Activity” that indicates the activation of a specific CA. Calculating the solution for each CA results in an overall cell (mRNA) profile, i.e. interrelated CA in terms of the simulated mRNA expressions. The higher the value of a PN-Activity, the higher the activity of a NP cell to express the respective mRNA. A PN-Activity of 0 means minimal (and not “zero”) expression of the respective mRNA.

### 2.3. Application of the PN-Methodology on the current system (Figure 1)

Functions that relate proteases and pro-inflammatory cytokines of NP cells to global stimuli (glucose, pH, magnitude, frequency) were used as previously discussed [21,22] (Appendix A). In contrast, all the data related to TIMP mRNA expressions was newly approximated and is subsequently introduced.

A data-driven approach was used to estimate the sensitivity of TIMP to glucose, pH, magnitude and frequency, as exemplified in Figure 2. Experimental data from Gilbert and colleagues were used to estimate the relationship between pH and TIMP [17] and data from Saggese et al. [20] was used to estimate the relationship between glucose and TIMP1. No data that directly related glucose concentrations with TIMP2 and TIMP3 was found. Therefore, a similar behavior as seen for TIMP1 was assumed. The response of TIMP to magnitude and frequency were obtained from Li et al. [9]. No information was found for TIMP2, hence, a similar behavior to TIMP1 was assumed. An overview of all the experimental data used is provided in Appendix A.

Functions that describe 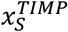 were fitted using the free online graphing calculator DESMOS (https://www.desmos.com/calculator). The parameters of a sigmoidal function were optimized to make the curve match the discrete data points from the experimental measurements.

To relate TIMP1 and TIMP2 mRNA expression to a physiological spectrum of magnitude, a generic function was used, as literature data reports the largest apparent mRNA fold-change between 0.8 and 1.3 MPa in accordance with the profile of the generic function. Generic functions are knowledge-based functions that were built on a variety of experimental data and reflected an overall range, within which a loading might be beneficial and in which range the loading might be detrimental. For magnitude, a shift from beneficial to detrimental (for persisting/chronic pressure values) is roughly estimated to be around 1 MPa (see Results, Figure 3; for more information [21], Appendix A).

Instructions on how to obtain weighting factors were previously provided [22]. In short: a maximum change in mRNA expression of cells exposed to different concentrations of stimuli (*ϵ*) was obtained from experimental research. To cope with the discrepancy between mRNA suppression (ranging from 0-1) and mRNA increase (ranging from 1<∞) a function of *ϵ* (*f*(*ϵ*)) was applied to turn mRNA suppression to semi bounded ranges by using reciprocal proportional relationships *f*(*ϵ*) = *ϵ* and 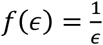 for ϵ >1 and for 0 < ϵ < 1, respectively. Thereby, *f*(*ϵ*) was understood as an estimation of the “cellular effort” to increase or decrease mRNA expressions compared to a control. To calculate the weighting factors 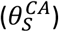, *f*(*ϵ*) for all significant relationships were normalized to the highest *f*(*ϵ*) within the network, whereas non-significant S-CA relationships were set to a 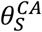 of 0.0010 to be less than 5% of the lowest experimentally determined weighting factor (here, the relationship between pH and TIMP1 with 0.0339, see Results, Table 2). The respective 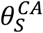 to estimate the effect of pro-inflammatory cytokines on TIMP were obtained from personal communication (Appendix A).

### 2.4 Predicting a pro-inflammatory environment

The protein levels of the pro-inflammatory cytokines, *Y*_*IL*1*B*_, *Y*_*TNF*_, were estimated based on their respective mRNA expressions (Eqs 3,4): PN-Activities of IL1B and TNF (*ω*_*IL*1*B*_, *ω*_*TNF*_) were normalized by their respective maximum possible mRNA expression (*ω*_*IL*1*B,max*_, *ω*_*TNF,max*_).

Maximum mRNA expressions were obtained by setting 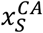 to a value of one under conditions that only activate the considered pro-inflammatory cytokine, i.e. when the inhibition term of the PN-Equation is zero (see [21]).

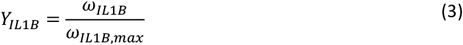

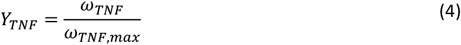

*Y*_*IL*1*B*_ and *Y*_*TNF*_ were then fed back into the intracellular network (Figure 1) as 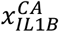 and 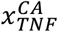, respectively (see Figure 2, example for 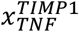) to calculate the mRNA expressions of TIMP1, TIMP2, TIMP3, MMP3 and ADAMTS4 of cells exposed to a pro-inflammatory environment.

### 2.5 Simulated conditions

PN-Activities for five daily human habits were simulated; “sitting”, “walking”, “hiking with 20kg extra weight”, “jogging” and “exposure to high vibration”. Human habits were chosen according to (i) prolonged activities of humans where cells might adapt to input stimuli, including possibly critical activities such as “exposure to high vibration” and “hiking with 20kg extra weight”. Intradiscal pressures were obtained from Wilke et al. [16]. Frequencies for walking and jogging were obtained from the literature [26,27]. A hiking pace of one step per second was assumed and a vibration frequency approximating an exposure to the vibration of an engine was reflected by 15.00 Hz.

All the human habits were simulated for three nutritional conditions that were obtained from mechano-transport Finite Element models [28] previously used as input for network modeling [22]. Two conditions simulated a non-degenerated IVD: one close to the Cartilage Endplates (“optimal” condition [24,29]); one around the mid-transverse plane of the IVD, in the anterior region of the NP, where the most adverse nutrient conditions were found within the mechanically loaded disc (“critical” condition). A third nutritional condition reflects an IVD also around the anterior mid-transverse plane of the NP, with a simulated early degenerated Cartilage Endplate [22] (Table 1).

**Table 1:**
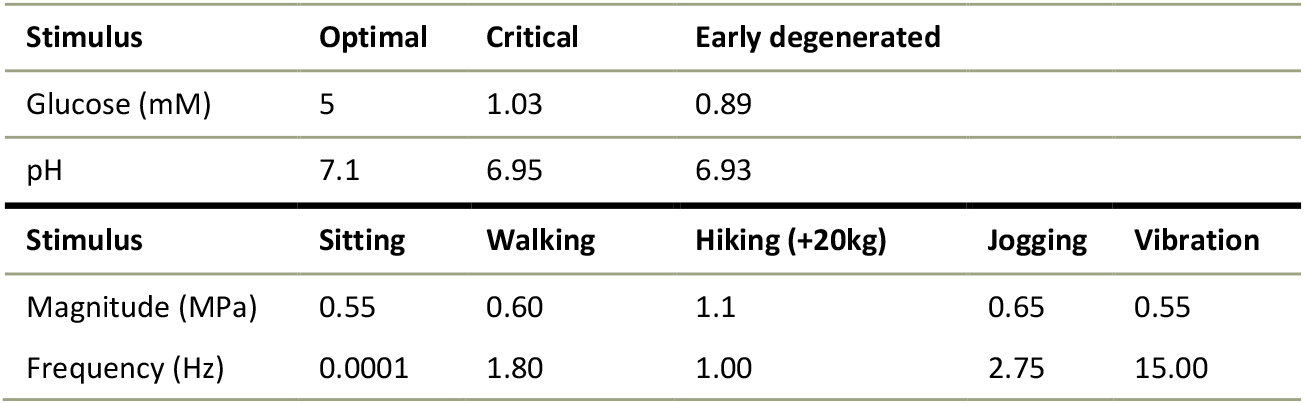
overview of stimulus values used to simulate five human habits within three nutrient environments.

| Stimulus | Optimal | Critical | Early degenerated |  |  |
| --- | --- | --- | --- | --- | --- |
| Glucose (mM) | 5 | 1.03 | 0.89 |  |  |
| pH | 7.1 | 6.95 | 6.93 |  |  |
| Stimulus | Sitting | Walking | Hiking (+20kg) | Jogging | Vibration |
| Magnitude (MPa) | 0.55 | 0.60 | 1.1 | 0.65 | 0.55 |
| Frequency (Hz) | 0.0001 | 1.80 | 1.00 | 2.75 | 15.00 |

**Table 2:**
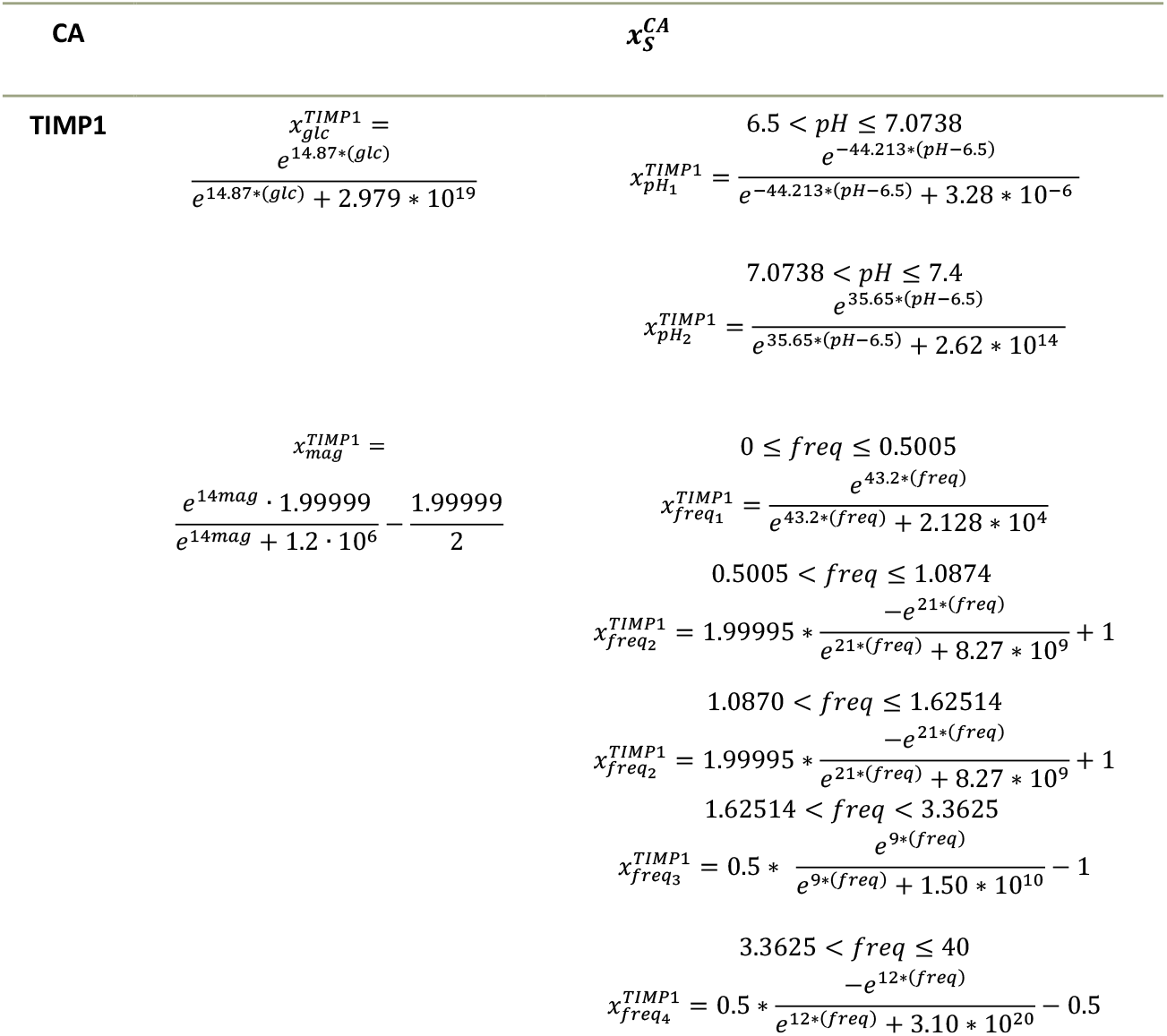
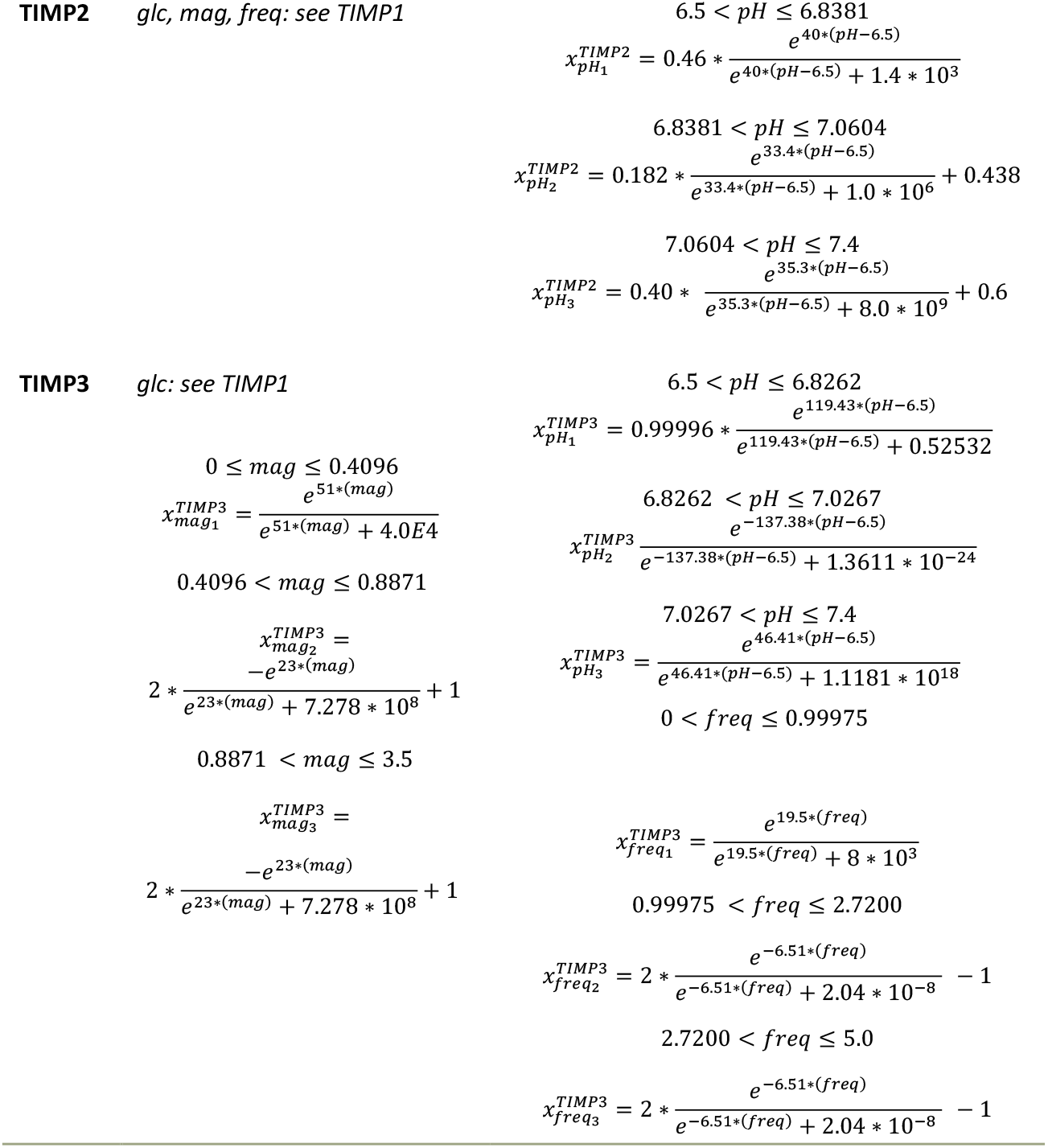
Functions that relate the Cell activity (CA) in terms of TIMP1, TIMP2 and TIMP3 mRNA expressions with varying physiological nutrient concentrations. Glucose (glc): 0< glc (mM) ≤ 5.0; pH: 6.5 < pH ≤7.4; magnitude (mag) 0 < mag (MPa)≤ 3.5; frequency (freq); 0 < freq (Hz) ≤ 40. If several (n) functions were required to describe a physiological range, functions were indicated with a subscript 1-n.

## 3 Results

### 3.1 System parameters

All the functions obtained to describe the response of TIMP to global stimuli are listed in Table 2.

A visualization of the responses of TIMP to magnitude and frequency is provided in Figure 3.

The weighting factors for this system are given in Table 3. The strongest S-CA relationship within the PN-System was found to be the relationship between pH and IL1B.

**Table 3:** Weighting factos for each cell activity (CA) and stimulus (S)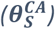. The highest weighting factor is shown in bold, weighting factors of non-significant S-CA relationships are set to 0.0010. S-CA relationships marked with * were not found in (or could not be directly translated from) literature and an approximate weighting factor was assumed. Mag: magnitude, Freq: frequency, Glc: glucose.

| Stimulus | mRNA | $\theta_S^{CA}$ | Stimulus | mRNA | $\theta_S^{CA}$ |
| --- | --- | --- | --- | --- | --- |
| glc | MMP3 | 0.0010 | mag | MMP3 | 0.0010 |
|  | ADAMTS4 | 0.0010 |  | ADAMTS4 | 0.0494 |
|  | IL1B | 0.0010 |  | IL1B | 0.0010* |
|  | TNF | 0.0010 |  | TNF | 0.0010* |
|  | TIMP1 | 0.0010 |  | TIMP1 | 0.0010 |
|  | TIMP2 | 0.0010* |  | TIMP2 | 0.0010* |
|  | TIMP3 | 0.0010* |  | TIMP3 | 0.0528 |
| pH | MMP3 | 0.3500 | freq | MMP3 | 0.1852 |
|  | ADAMTS4 | 0.0704 |  | ADAMTS4 | 0.0988 |
|  | IL1B | **1.0000** |  | IL1B | 0.0010* |
|  | TNF | 0.0010 |  | TNF | 0.0010* |
|  | TIMP1 | 0.0339 |  | TIMP1 | 0.0353 |
|  | TIMP2 | 0.0417 |  | TIMP2 | 0.0353* |
|  | TIMP3 | 0.0010 |  | TIMP3 | 0.0523 |
| IL1B | MMP3 | 0.1333 | TNF | MMP3 | 0.3310 |
|  | ADAMTS4 | 0.0010 |  | ADAMTS4 | 0.0710 |
|  | TIMP1 | 0.0010* |  | TIMP1 | 0.0010* |
|  | TIMP2 | 0.0010* |  | TIMP2 | 0.0010* |
|  | TIMP3 | 0.0010 |  | TIMP3 | 0.0010 |

### 3.2. System predictions

Cells not exposed to pro-inflammatory cytokines showed a high anabolic CA with elevated TIMP2 and TIMP3 mRNA expressions in all human habits throughout all nutrient conditions. TIMP1 only showed notable upregulation in “hiking with 20 kg extra weight”. Elevated protease mRNA expression was only predicted for hiking with additional weight and vibration, whilst remaining minimal for any other condition. Exposure to pro-inflammatory cytokines had a marginal effect on TIMP but caused an upregulation of proteases in all the simulated conditions. Upregulation of MMP3 was more pronounced than of ADAMTS4 (Figure 4) and notably upregulated towards more adverse nutrient conditions.

**Figure 4:**
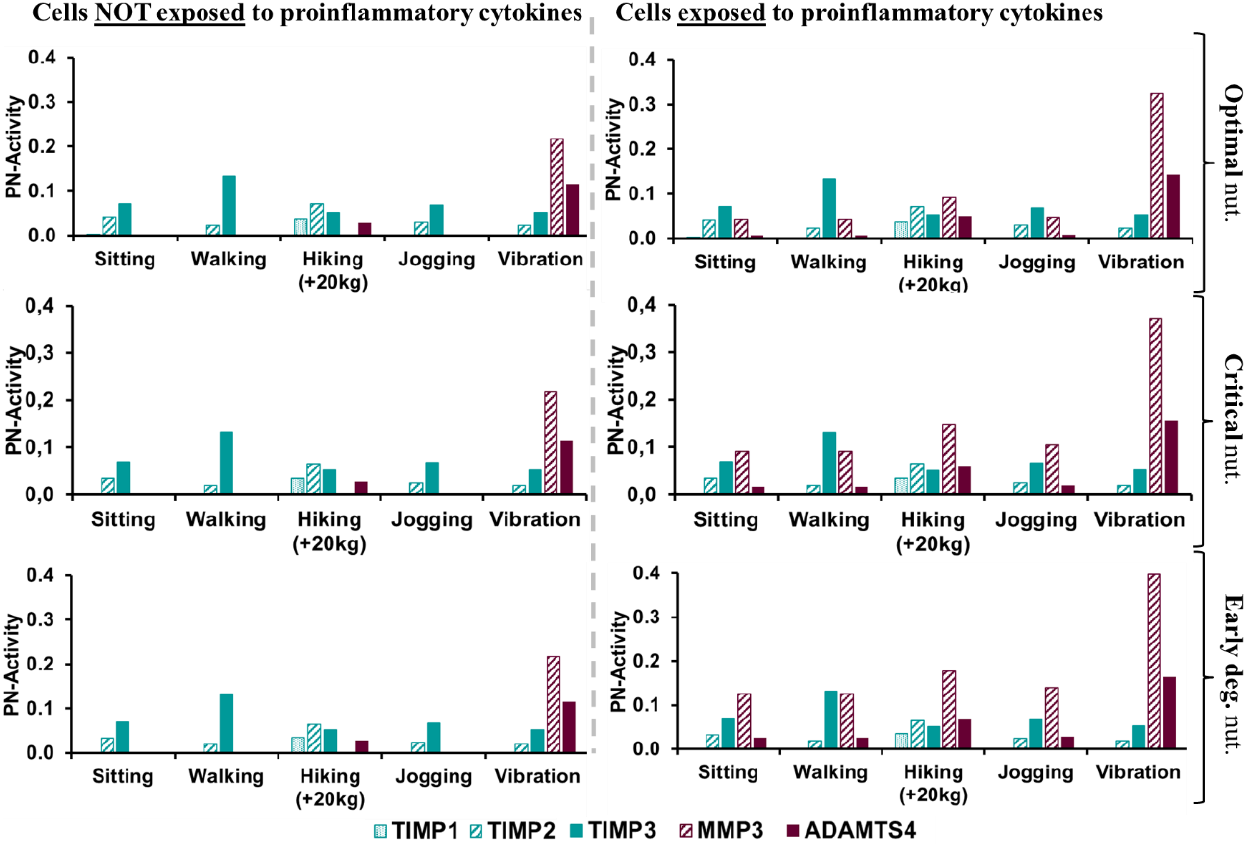
Predicted mRNA profiles for cells not exposed and exposed to pro-inflammatory cytokines for five human habits: sitting, walking, hiking with 20kg extra weight, jogging and exposure to high vibration, and within three different nutrient (nut.) environments being optimal (pH 7.1, 5 mM glucose), critical (pH 6.95, 1.03mM glucose) and early degenerated (pH 6.93, 0.89 mM glucose).

In terms of pro-inflammatory cytokines, the mRNA expression of IL1B was predicted to be higher than the mRNA expression of TNF. PN-Activities of both, IL1B and TNF slightly rose under more adverse nutrient concentrations (Figure 5, left). IL1B protein synthesis was overall constantly low throughout different nutrient and loading conditions, whilst TNF largely increased with adverse nutrient and loading conditions (Figure 5, right).

**Figure 5:**
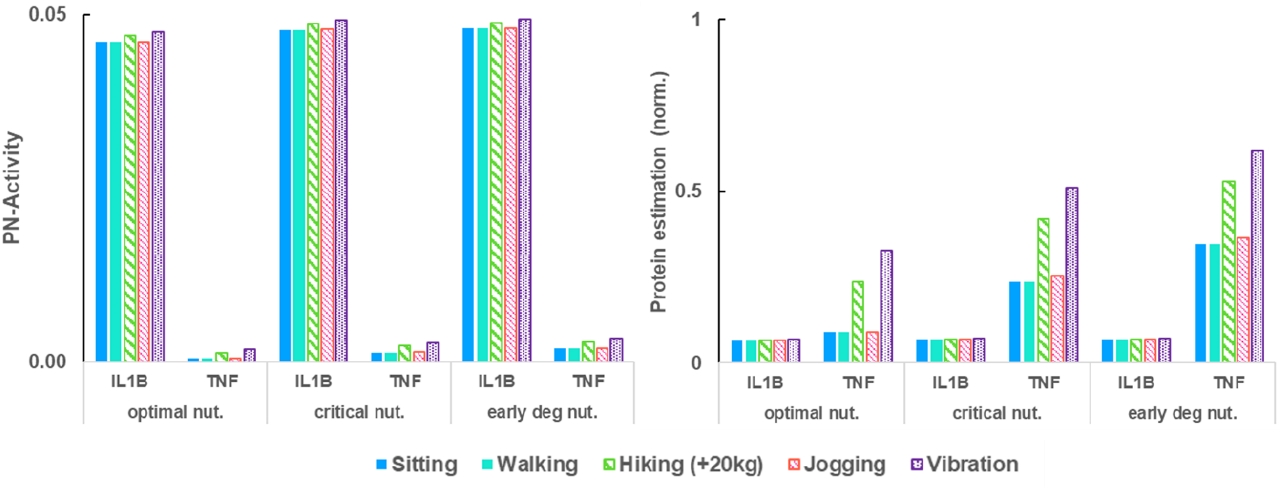
mRNA expressions (left) and protein estimations (right) of TNF and IL1B under different nutritional (nut.) and loading conditions.

## 4 Discussion

### 4.1 TIMP and protease cell profiles (Figure 4)

Unique TIMP and protease CA for a variety of human habits for three different nutrient concentrations were obtained for NP cells exposed and not exposed to proinflammatory cytokines. Such cell profiles provided by the PN-Methodology might be read as relative CA of one (average) cell, or it can be interpreted as the average response of various cells. This becomes particularly important when comparing simulation results with experimental data of the percentage of immunopositive cells for TIMP and proteases for different stages of degeneration [5], and with evaluations of the effect of a combined IL1B and TNF enriched culture medium on bovine NP biopsies [30].

With more adverse nutrient concentrations, a catabolic shift was predicted, which is particularly pronounced in cells exposed to pro-inflammatory cytokines. Low effect of varying physiological and early degenerated nutrient concentrations within short timeframes agrees with general expectations that degenerative processes take months to years to evolve. However, literature shows that with rising degenerative levels, also the relative percentage of cells immunopositive for pro-inflammatory cytokines rise [11,21], hence, apart from the individual catabolic shift of one cell also the number of cells rise that show a catabolic activity.

Regarding specific human movement habits, a strong catabolic shift due to an exposure to vibration (15 Hz, Table 1) is in agreement with findings reporting that a low level of frequency is required to maintain the functionality of the IVD [9,31,32]. However, high vibration might often be damped, so that detrimental loading might not reach the NP cell micro-environment. Interestingly, an elevated TIMP1 expression was only predicted by the model for hiking with extra weight, the only condition that exceeds intradiscal loading pressure of 1 MPa (Table 1). This was supported by experimental findings that TIMP1 expressions in bovine NP tend to be upregulated under loading conditions exceeding 1MPa [33].

As per pro-inflammatory cytokine exposure, the model predicts an overall catabolic shift (Figure 4), which is supported by various experiments [11,22,34,35]. Thereby, the catabolic shift is characterized by a stagnant TIMP mRNA expression and a rise of proteases. Similar findings were provided for TIMP3, with a stagnant percentage of immunopositive cells, while the percentage of immunopositive cells of any other of the mRNA expressions tackled in this study significantly rose throughout degenerative stages [5]. A stagnant prediction of TIMP1 and TIMP2 is attributed to the lack of data about the relationship between IL1B and TNF on TIMP1 and TIMP2, which required the use of data of TIMP3 also for TIMP1 and TIMP2 (see also Appendix A, C). As soon as experimental data about the individual effect of IL1B and TNF on TIMP1 and TIMP2 is available, the PN-modelling design allows for a smooth implementation of this information.

Regarding proteases, a rise of MMP3 for all the human activities due to presence of pro-inflammatory cytokines was predicted, which is in concordance with a positive correlation between increasing percentages of cells immunopositive for MMP3, degeneration grades [5] and correspondingly to combined IL1B and TNF exposure [30].

ADAMTS4 mRNA expression was predicted to be minimal and low, without and with pro-inflammatory cytokine exposure, respectively. This was confirmed by a non-significant change in ADAMTS4 expression due to combined IL1B and TNF exposure [30]. ADAMTS4 levels were, however, always increased under hiking with extra weight and with exposure to high vibration (Figure 4). This confirms that such loading conditions might contribute to IDD by reflecting detrimental effects as found in weight lifting, heavy work (carpenters) [36,37] and whole body vibration (machine drivers) [37], respectively. Hence, results support the benefit of hip-belts while hiking with extra load.

In terms of relative mRNA predictions within the cell profiles; TIMP3 mRNA expression was predicted by the model to be the overall most upregulated TIMP mRNA expression, independently from the presence of pro-inflammatory cytokines. This goes along with findings of Le Maitre et al., 2004 [5], who observed that throughout degeneration TIMP3 immunopositivity was always highest compared to TIMP1 and TIMP2 immunopositivity. Thereby, a rise of TIMP2 was only found for importantly degenerated IVD, which goes along with our findings that initially only encompass early degenerative states (see 4.2 for relativization). However, a rise of TIMP1 immunopositivity was already observed at lower degenerative states [5], which might suggest a underprediction of TIMP1 by the model at early-degenerative nutrient concentration. Conversely, other experimental studies regarding the effect of TIMP1 on TNF and IL1B on bovine whole organ cultures confirmed non-significant alterations of TIMP1 within combined TNF and IL1B exposure for up to seven days of exposure, before becoming significant [30]. Such findings emphasize the importance of time-sensitivity to duly categorize and understand CA in both, experimental and *in silico* research. While the effect of long-term exposure is yet to be fully investigated for the current TIMP and protease regulations, the PN-Methodology, was designed for the continuous integration of time-sensitive approximations [21], and can, thus, eventually provide more insight into temporal considerations.

In terms of proteases, MMP3 immunopositivity was predicted by the model to be minimal in absence of pro-inflammatory cytokines for all non-catabolic loading conditions. This is aligned with negligible MMP3 immunopositivity in non-degenerated NP cells [5]. However, the overall low ADAMTS4 mRNA expression stood in contrast by findings of Le Maitre and colleagues [5], who found that around 20% of NP cells expressed ADAMTS4 already at a non-degenerative state which then rises throughout degeneration. This indicates that changes of ADAMTS4 might be correctly predicted by the model, whilst the overall PN-Activity compared to other mRNA expressions might be underpredicted. Reasons therefore might be important discrepancies between the used experimental data (Appendix A) and actual human *in vivo* expressions or the neglection of an additional (global) stimulus that strongly regulates ADAMTS4 mRNA expression in addition to the four tackled global stimuli. This outcome shows the potential of the PN-approach to identify if key relevant stimuli were considered for a certain mRNA expression.

Hence, the direct use of experimental data due to the PN-modeling approach allows to maintain biological proximity is beneficial, but limitations regarding data availability and non-standardized experimental protocols need to be carefully tackled. With this regard, effort was made to choose studies where less differences in the experimental setup were found. In cases where experimental data was only found for either TIMP1 or TIMP2, the TIMP type that lacked relevant data was approximated to the results found for the other. Eventually, limitations of the current validation should be highlighted, such as assuming a direct relationship between more mRNA expression and a higher percentage of immunopositive cells, even though, theoretically, higher amounts of mRNA expression could also arise from a higher mRNA productivity of each cell, without upregulating the actual amount of immunopositive cells. With this regard, the current validations were based on both, percentages of immunopositive cells [5] and effective mRNA expressions due to combined IL1B and TNF exposure [30].

To evaluate the inhibitory potential of TIMP on proteases arising from the presented cell profiles, further analysis was conducted (Appendix B), predicting TIMP to efficiently downregulate proteases in all conditions but exposure vibration in cells not exposed to pro-inflammatory cytokines. Due to pro-inflammatory cytokine exposure, particularly the inhibition of MMP3 was predicted to be impeded.

### 4.2 The pro-inflammatory environment (Figure 5)

At the mRNA level, both IL1B and TNF were predicted to have a relatively low expression with a PN-Activity of less than 0.05, thereby, IL1B being higher expressed than TNF. This tendency was also found in experimental research, where both the percentage of cells that express IL1B and its receptor tended to be higher than the percentage of cells that express TNF and its receptor [11]. Interestingly, a translation to proteins based on the approach of the PN-Methodology indicates that the proinflammatory environment causing the catabolic shift in the cell profiles shown in Figure 5 is rather provoked by an increase of TNF, than of IL1B proteins. This finding is biologically interesting, related to the generally assumed detrimental effect of TNF [13], in contrast to a possible physiological role of IL1B at low concentrations [38]. Given that the PN-Methodology only describes relative changes of a CA, a baseline of pro-inflammatory cytokine levels under physiological concentrations would be required to further evaluate model predictions. In other words, measurements would be required that contrast quantitative IL1B and TNF protein levels under non-degenerated and early degenerated conditions. In terms of quantity, however, only few studies actually investigated physiological cytokine expression within IVD tissue [39–41].

Importantly, those studies suggest that pro-inflammatory cytokine levels range within pg/ml, whereas most experimental studies focusing on the effect of pro-inflammatory cytokines, including the ones used for this study (Appendix A), stimulate cell cultures with around 10ng/ml, which seems to be largely hyper-physiological. In contrast, experimental studies simulating mechanical or nutritional environments that are often stimulated under physiological conditions (see e.g. studies listed in Appendix A). This bias is translated to in silico research and, consequently, the hereby presented model predictions of the pro-inflammatory status might be biased, or possibly reflecting higher degenerative states in terms of pro-inflammatory environments than early degeneration.

Further discussion focused on mathematical aspects of the PN-Methodology and the impact of uncertainties due to a lack of experimental data are provided in Appendix C.

## 5 Conclusions

This work aimed to provide insight into the dynamics of TIMP, proteases and pro-inflammatory cytokines in response to the multifactorial NP environment. Approximations were obtained through the PN-Methodology, that was used (i) to estimate relative mRNA expressions of TIMP and proteases and (ii) to approximate a proinflammatory environment. The strength of the PN-Methodology is its efficiency to comprehensively estimate CA in complex multifactorial environments, without the need to simulate intracellular regulatory networks. Moreover, with the data directly derived from experimental research, an approximation from the mRNA to protein level was deduced. Predicted cell responses largely agree with independent experimental studies, indicating that the PN-Methodology proves utility in estimating CA in response to complex stimulus environments. Further work should investigate the generalization of this methodology towards other tissues and its validity to link mRNA expressions with corresponding protein synthesis.

## AUTHOR CONTRIBUTIONS

Study concept and design: Baumgartner L., Noailly J. Acquisition of material and data: Witta S., Baumgartner L. Data analysis: Witta S., Baumgartner L. Preparation of the manuscript: Baumgartner L. Critical reviewing and approval of the manuscript: all authors.

## ACKNOWLEDGEMENTS

The authors thank Paola Bermudez for providing information about her cell culture experiments within the framework of disc4all [Disc4All-MSCA-2020-ITN-ETN 955735] regarding the response of TIMP3 to IL1B and TNF enriched culture media.

This work was financially supported by the European Research Council [ERC-2021-CoG-O-Health-101044828].

## CONFLICT OF INTEREST STATEMENT

The authors declare no conflict of interest.

## DATA AVAILABILITY STATEMENT

All the functions 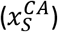 and values of the weighting factors 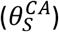 are provided within this work or within previous open access publications. The methodology on how to build PN-Equations is provided within this work and more extensively in previous work [21]. Additionally, we are working on curating the full code for online redistribution, which shall be ready by the time of the review.

## Notes

### Competing Interest Statement

The authors have declared no competing interest.

