## Appendix for "Parallel networks to predict TIMP and protease cell activity of Nucleus Pulposus cells exposed and not exposed to pro-inflammatory cytokines": Baumgartner_et_al_2024_TIMP_proteases_appendix.pdf

### Appendix A

The compilation of experimental studies to characterize the variable $x$ of the C-SA relationships is listed in table A1.

Table A1: overview of the experimental literature used to create the $x$ values of the S-CA relationships. Note that no x-values are required for pro-inflammatory cytokines as they were directly created by the model (see main study).

| Stimulus | mRNA | Cell type | Experimental source | Functions for x provided in |
|---|---|---|---|---|
| Glc | MMP3 | Human | [1] | [2] (supplementary material) |
| | ADAMTS4, IL1B, TNF | Bovine | [2] | [2] (supplementary material) |
| | TIMP1 | Human | [3] | This study |
| | TIMP2 | *No information* | | *TIMP1-like nature assumed* |
| | TIMP3 | *No information* | | *TIMP1-like nature assumed* |
| pH | MMP3, ADAMTS4, IL1B, TNF, TIMP1, TIMP2, TIMP3 | Human | [4] | [2] (supplementary material) |
| Magnitude | MMP3, ADAMTS4 | Knowledge-based, based on several studies | | [5] (generic function) |
| | IL1B | Generic behavior, same as MMP3, ADAMTS4 | | [5](generic function) |
| | TNF | Generic behavior, same as MMP3, ADAMTS4 | | [5](generic function) |
| | TIMP1 | Porcine | [6] | [5](generic function) |
| | TIMP2 | *No information* | | [5](generic function) |
| | TIMP3 | Porcine | [6] | This study |
| Frequency | MMP3, ADAMTS4 | Knowledge-based, based on several studies | | [5] (generic function) |
| | IL1B, TNF | *No information* | | [5] (generic function) |
| | TIMP1, TIMP3 | Porcine | [6] | This study |
| | TIMP2 | *No information* | | *TIMP1-like nature assumed* |

The generic function mentioned in Table A1 refers to a function covering physiological ranges of loading mag and freq that was approximated to reflect a general anabolic and catabolic behavior, respectively. Generic functions are a knowledge-based approximation in case of data-based experimental information was not available. The generic functions were previously published and explained in-depth [5] and assume a change from overall anabolic to catabolic effects around 1MPa of magnitude and 3Hz of frequency, respectively. Generic functions were applied to approximate the behavior of proteases exposed to loading parameters and to describe the response of TIMP1 and TIMP2 to magnitude.

Weighting factors ($\theta$) depend on the actual PN-System and are added for the current system (Table A2).

| Table A2: Compilation of the capacities ($\epsilon$), cellular efforts $f(\epsilon)$, and calculated weighting factors $\theta_S^{CA}$, characterizing the S-CA relationships of the 1st and 2nd level CA of Fig (1). NS: not significant, *: no information found | | | | | | |
|---|---|---|---|---|---|---|
| Stimulus | mRNA | $\boldsymbol{\epsilon}$ | $\boldsymbol{f(\epsilon)}$ | $\boldsymbol{\theta_S^{CA}}$ | Source | Cell Type |
| Glc | MMP3 | NS | - | 0.00100 | [1] | Human |
| | ADAMTS4 | NS | - | 0.00100 | [2] | Bovine |
| | IL1B | NS | - | 0.00100 | [2] | Bovine |
| | TNF | NS | - | 0.00100 | [2] | Bovine |
| | TIMP1 | NS | - | 0.00100 | [3] | Human |
| | TIMP2 | NS | - | 0.00100 | * (*TIMP1-like impact assumed)* | Human |
| | TIMP3 | NS | - | 0.00100 | * (*TIMP1-like impact assumed)* | - |
| pH | MMP3 | 28.7 | 28.7 | 0.35 | [4] | Human |
| | ADAMTS4 | 5.7 | 5.7 | 0.0704 | [4] | Human |
| | IL1B | 81.000 | 81.000 | 1.0000 | [4] | Human |
| | TNF | NS | - | 0.0010 | [4] | Human |
| | TIMP1 | 2.742 | 2.742 | 0.0339 | [4] | Human |
| | TIMP2 | 0.296 | 3.379 | 0.0417 | [4] | Human |
| | TIMP3 | NS | - | 0.0010 | [4] | Human |
| Mag | MMP3 | NS | - | 0.0010 | [7] | Rat |
| | ADAMTS4 | 4 | 4 | 0.0494 | [7] | Rat |
| | IL1B | NS | - | 0.0010 | [8,9](Indications) | |
| | TNF | NS | - | 0.0010 | [10] (Indications) | |
| | TIMP1 | NS | - | 0.0010 | [6] | Porcine |
| | TIMP2 | NS | - | 0.0010 | * (*TIMP1-like impact assumed)* | - |
| | TIMP3 | 4.2731 | 4.2731 | 0.0528 | [6] | Porcine |
| Freq | MMP3 | 15 | 14.7059 | 0.1852 | [7] | Rat |
| | ADAMTS4 | 8 | 3.0075 | 0.0988 | [7] | Rat |
| | IL1B | NS | - | 0.0010 | * (discussed in [5]) | - |
| | TNF | NS | - | 0.0010 | * (discussed in [5]) | - |
| | TIMP1 | 2.8571 | 2.8571 | 0.0353 | [6] | Porcine |
| | TIMP2 | 2.8571 | 2.8571 | 0.0353 | * (*TIMP1-like impact assumed)* | - |
| | TIMP3 | 4.2353 | 4.2353 | 0.0523 | [6] | Porcine |
| IL1B | MMP3 | 10.8 | 10.8 | 0.1333 | [11] | Human |
| | ADAMTS4 | NS | - | 0.0010 | [11] | Human |
| | TIMP1 | NS | - | 0.0010 | * (*TIMP3-like impact assumed)* | - |
| | TIMP2 | NS | - | 0.0010 | * (*TIMP3-like impact assumed)* | - |
| | TIMP3 | NS | - | 0.0010 | Personal communication out of the | Bovine |

| | | | | | | |
|---|---|---|---|---|---|---|
| | | | | | disc4all consortium [Disc4All-MSCA-2020-ITN-ETN 955735] | |
| TNF | MMP3 | 26.85 | 26.85 | 0.331 | [2] | Bovine |
| | ADAMTS4 | 5.77 | 5.77 | 0.071 | [2] | Bovine |
| | TIMP1 | NS | - | 0.0010 | * (*TIMP3-like impact assumed)* | Bovine |
| | TIMP2 | NS | - | 0.0010 | * (*TIMP3-like impact assumed)* | - |
| | TIMP3 | NS | - | 0.0010 | Personal communication, out of the disc4all consortium [Disc4All-MSCA-2020-ITN-ETN 955735] | Bovine |

In-house experiments were done on bovine NP cells cultured in alginate beads. Beads were either exposed to 10 ng/ml TNF or to 10ng/ml of IL1B and compared to a control without pro-inflammatory cytokine exposure. It was found that the mRNA expression of TIMP3 normalized to encapsulation day was not significantly altered after one week of cytokine exposure. No information about the effect of IL1B and TNF on TIMP1 and TIMP2 was found, which led to an initial assumption of a non-significant impact also for those mRNA expressions.

### Appendix B

#### Analysis of residual proteins within the tissue

To estimate protein levels of TIMP1, TIMP2, TIMP3, MMP3 and ADAMTS4 under presence and absence of pro-inflammatory cytokines a linear relationship between mRNA expression and corresponding protein synthesis was assumed. For the interaction between proteases and TIMP, the binding capacity of proteins such as the stochiometric ratio was considered. Hence, the modeling pipeline was extended by combining it with a conventional directed network modeling approach (Figure B1).

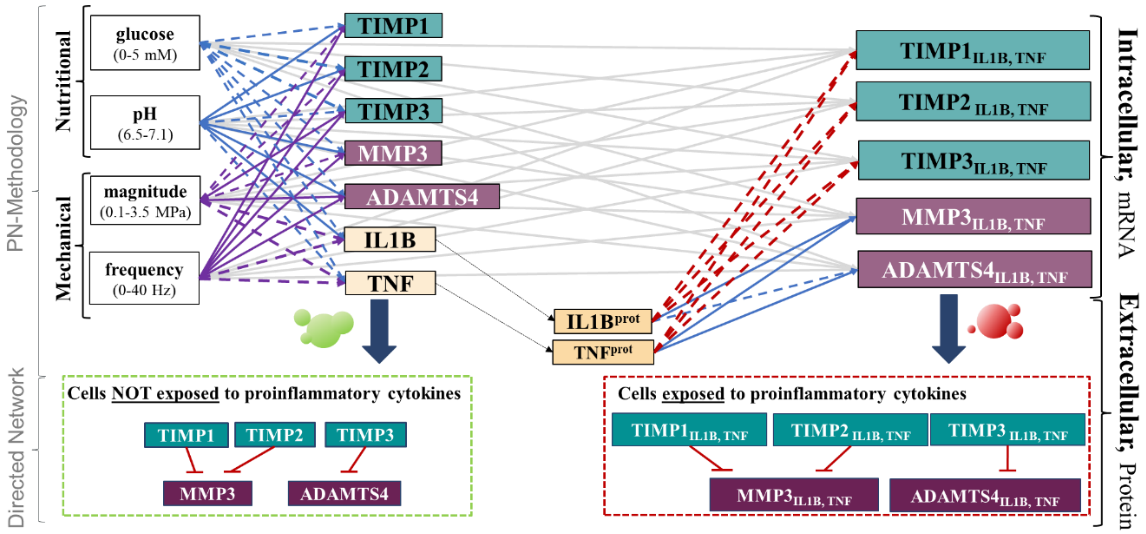


*Figure B1: Overview of the coupled modeling approach. Intracellular: network as pesented in Figure 1 of the main text. Extracellular: a directed network modeling approach was used to estimate the presence of TIMP proteins and proteases within the Nucleus Pulposus extracellular matrix.*

The binding capacity of TIMP proteins on proteases was estimated through the inhibitory constant ($K_i$) (Table B1).

*Table B1. Inhibitory constant values of TIMP1 and TIMP3 for the proteases MMP3 and ADAMTS4 given in nanomolar. NS: not significant, $\gamma$: normalized strength of the affinity.*

| Inhibitor | $K_{i,MMP3}$ | $\gamma_{MMP3}$ | $K_{i,ADAMTS4}$ | $\gamma_{ADAMTS4}$ | Reference |
|---|---|---|---|---|---|
| TIMP1 | 1.9 ± 0.1 nM | 1 | NS | 0.001 | [12] |
| TIMP3 | 66.9 nM | 0.028<br>(=1.9/66.9) | 3.3 nM | 1 | [13] |

Given that the affinity of TIMP3 reflects only 2.8%, (Table B1) of the affinity of TIMP1 to bind MMP3, TIMP3 was not considered as a relevant inhibitor of MMP3. A similar affinity for TIMP1 and TIMP2 for ADAMTS4 was assumed based on key structural similarities in the N-terminal domain for TIMP1 and TIMP2 [14]. Hence, directed networks were built by assuming an inhibition of TIMP1 and TIMP2 on MMP3 and an inhibition of ADAMTS4 due to TIMP3 (Figure B1, extracellular).

As for the stochiometric ratio, it was found that TIMP interact with the active site cleft of ADAMTS and MMP through the formation of a tightly bound complex in a 1:1 stoichiometric ratio, i.e., one TIMP protein is able to inhibit one protease [15].

First approximations of the relative amount of TIMP and proteases within the extracellular matrix were individually analyzed for each human habit. Estimations were done assuming optimal binding conditions where all proteins are available to interact, without additional estimations of possible spatiotemporal effects. Therefore, mRNA profiles of respective binding partners, i.e. MMP3-TIMP1,2 and ADAMTS4 – TIMP3 for each condition were individually translated to estimate the residual protein abundancy after the binding process. Thereby normalized protein levels ($Y_{TIMP1}, Y_{TIMP2}, Y_{TIMP3}, Y_{MMP3}, Y_{ADAMTS4}$) were obtained by setting the higher expressed binding partner to one and the less expressed binding partner to its relative normalized value, according to the estimated mRNA levels. Additive effects in case of TIMP1 an TIMP2 were assumed, leading to an estimated residual (subscript "res") potein levels as given in Eq. B1 and Eq. B2:

$$Y_{res,MMP3/TIMP1,2} = |Y_{MMP3} - (Y_{TIMP1} + Y_{TIMP2})| \quad \text{(B1)}$$
$$Y_{res,ADAMTS4/TIMP3} = |Y_{ADAMTS4} - Y_{TIMP3}| \quad \text{(B2)}$$

### Results

MMP3 and ADAMTS4 were predicted to be efficiently bound in absence of pro-inflammatory cytokines, with almost 100% of residual TIMP in the ECM. This was true for all human habits, but exposure to vibration, where around 80% of MMP3 and around 50% of ADAMTS4 were predicted to reside within the ECM after an optimal binding process. In presence of pro-inflammatory cytokines, the binding of MMP3 was predicted to be worse than that of ADAMTS4, with rising catabolism under adverse nutrient conditions (Figure B2).

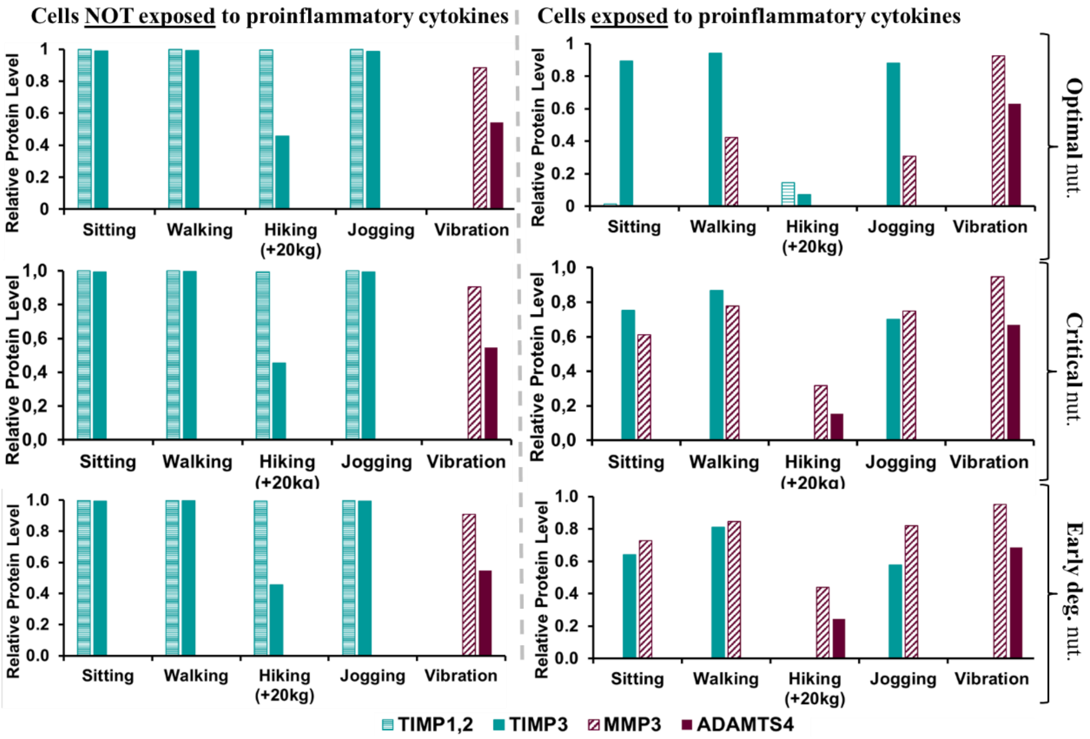


*Figure B2: Predicted normalized residual proteins for cells not exposed and exposed to pro-inflammatory cytokines for five human habits: sitting, walking, hiking with 20kg extra weight, jogging and exposure to high vibration, and within three different nutrient environments; optimal (pH 7.1, 5 mM glc), critical (pH 6.95, 1.03mM glc) and early degenerated (pH 6.93, 0.89 mM glc).*

### Discussion

At a protein level model results suggest that proteases could be strongly downregulated by TIMP in all cases but in case of exposure to high vibration. Thereby, a high overshoot of TIMP proteins (nearly 100%) after binding was predicted in most human activities. Findings agree with Molinos et al., 2015 [16], who report low expressions of MMP3 and ADAMTS4 in a non-degenerated NP. An overshoot of TIMP is comprehensible in the light of both, further roles of TIMP within the NP, e.g. the inhibition of the activity of TNF converting enzyme and the suppression of vascular ingrowth via its binding to VEGF through TIMP3 [17] and the importance of spatiotemporal effects of proteins within the extracellular matrix. Hence, spatial proximity between proteases and their inhibitors were biologically required within the lifespan of proteins, which possibly requires an

important amount of TIMP proteins compared to protease levels. In this work, spatiotemporal effects were not yet considered. An integration could be achieved by a coupling of the current numerical model to an Agent-based approach. Such information could then retroactively be incorporated into the current model to improve estimations of protein availability.
Due to pro-inflammatory cytokine exposure (Figure B2, right) inhibition of especially MMP3 failed. Catabolism got generally more severe under progressive nutrient deprivation.

To predict inhibition, a selective binding of TIMP1,2 on MMP3 and TIMP3 on ADAMTS4 was assumed, which is based on differences of the TIMP inhibitory activity of its N-terminal domain [12]. However, it can be assumed that a low inhibiting effect of TIMP3 on MMP3 might additionally take place. Therefore, the current study might underestimate actual binding capacities of MMP3. Eventually, a linear relationship was assumed between semiquantitative mRNA and protein levels for those predictions, which, therefore, might only reflect an overall tendency.

### Appendix C

#### Mathematical aspects of the methodology

As compared to the previously explained PN-Framework [5], in this system, theta values required a broadening of their range from the initially introduced $0.01 \leq \theta_S^{CA} \leq 1$ to $0.001 \leq \theta_S^{CA} \leq 1$ due to larger differences in mRNA expression fold changes characterizing S-CA relationships. This does not imply any numerical limitations given that the PN-Methodology technically allows theta values equal or bigger than 0.0001.

Moreover, as opposed to previously published work [5], in the current approach mRNA expressions of pro-inflammatory cytokines were calculated relative to the mRNA expressions of TIMP and proteases. This allowed to estimate a range of pro-inflammatory cytokine expression relative to any other mRNA and use the individual maximal mRNA expressions as a reference to approximate relative protein levels (Eqs 3, 4, main manuscript). As a result, important increases of protein levels based on marginal variations of mRNA expressions were predicted by the PN-Methodology, the biological interpretability of which was discussed (section 4.2, main manuscript). Given that such predictions strongly rely on adequate experimental data, a lack thereof might very much influence such findings.

For the current system, uncertainties due to a lack of experimental data are reflected in Figure C1.

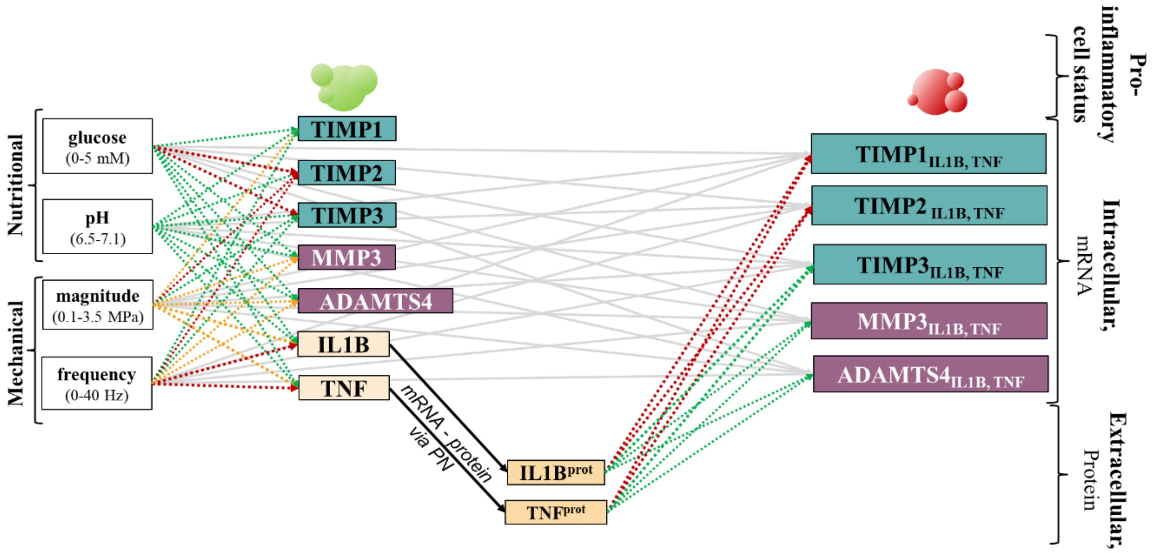


*Figure C1: overview of the biological quality of the S-CA relationships. Whilst most relationships were directly derived from experimental measurements (green), seven relationships were approximated by a generic behavior (orange), and for ten relationships a similar behavior as for closely related proteins was assumed (red).*

There was a lack of data-based experimental information in 17 S-CA relationships, out of which 7 S-CA relationships were approximated by knowledge-based generic behavior and for 10 S-CA relationships a similar behavior as seen for closely related proteins was assumed.

In case of TIMP subfamilies, a similar behavior among them was expected, thereby, a closer relationship between TIMP1 and TIMP2 was assumed due to their similarity in terms of protease binding (Appendix B). The effect of freq on IL1B and TNF was previously discussed [5]: a non-significant effect of freq on pro-inflammatory cytokines was assumed and mRNA expression followed a generic behavior.

**References Appendix**
